# Stimulus-dependent depth constancy during head tilt

**DOI:** 10.1101/2022.03.04.483050

**Authors:** Jan Skerswetat, Andrea Caoli, Peter Bex

## Abstract

Stereopsis is traditionally measured with noise-based stereo tests while the observer views the test in primary gaze. We investigated the effect of stimulus sparseness and axial variations of interocular disparity induced via head rotations.

First, we measured stereoacuity using a 4-Alternative-Forced-Choice (4-AFC) task with three uncrossed and one crossed disparity bandpass-filtered circles on a passive-3-D-monitor. Ten binocularly-normal adults fixated a central cross and clicked on the circle withcrossed disparity for forty trials/condition. Observers adopted head tilts of 0° or ±20° pitch, roll, or yaw, enforced with an innertial measurement unit and fixation enforced with an eye tracker. Next, we measured stereoacuity in 8 adults while either the head (H), monitor (M), or both (B) were tilted 0°, ±22.5°, or ±45° roll in random order (eighty trials/condition) using a 4-AFC task and random-dot stimuli. Head tilts did not signifcantly alter stereoacuity using narrow-band stimuli(p>0.05), despite that IPDs and the axis of disparity were differentially affected by the tilts. However, for random dot stimuli, stereoacuity decreased with increasing orientation difference between the head and monitor (H and M: p<0.05; B: p>0.05].

Head tilt decreases IPD and rotates the axis of interocular disparity, however, these manipulations affect stereoacuity when measured with noise stimuli but not with sparse stimuli. The results are consistent with a vestibular input to stereoscopic disparity processing that can be detected by sparse stimuli but is masked by dense stimuli. The results have implications for natural vision and for clinical screening in patients with abnormal head posture.

**Significance statement:** Depth perception is a critical feature of human vision and it is thought that the ability to perceive stereoscopic depth is bound to an essentially eye-fixed, horizontal disparity of each image that rapidly deteriorates away from that limited horizontal axis. In a set of head tilt experiments, we varied the orientations of stereoscopic images and demonstrate that stereoacuity remains constant when deploying sparse narrow-band stimuli, and only worsens when using fine-detailed noise stimuli that mask off-axis disparities. These results shine new light upon the debate of neuroplasticity of stereo vision. Moreover, the results are consequential for diagnosis and treatment in people with atypical head- and eye alignment, such as for patients with torticollis or strabismus.

## 1. Introduction

Depth perception is a hallmark feature of binocular visual systems in a variety of species (1) and for humans, it plays a key role in activities of daily living including occupational and recreational tasks. Clinical stereo tests are carried out while the patient views the test upright, in approximately primary gaze, at reading distance, while holding an upright head posture. Outside of the testing setting, patients will however hold head tilts to the side, e.g. when crossing a street (i.e. horizontal rotation *or yaw*), when looking up to see a bird in a tree (i.e. vertical rotation or *pitch*), or tilting sideward e.g. when looking underneath a table for a power outlet (i.e. torsional rotation or *roll*). Increasing the head tilt therefore increases the difference between head tilt and counter roll and thus increases the rotation of the retinal images relative to that during the horizonal upright head position.

Head yaw and roll change interpupilary distance (IPD) by the simple cosine of the yaw or roll angle θ, relative to primary gaze 0°:

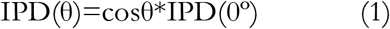

For small depths (*d*), at viewing distance *D*, retinal disparity (*R*) is affected by IPD approximately as:

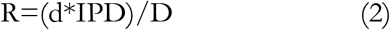

Thus, retinal diparity decreases in direct proportion to IPD as yaw and roll increase. Since IPD affects stereoacuity (2), head tilt should directly affect stereoacuity. Furthermore, head roll also introduces a vertical offset between the corresponding visual axes of the eyes, which should disrupt the horizontal arhitecture of binocular correspondence fields (Cumming, 2002; Cumming & DeAngelis, 2001; Read & Cumming, 2017). Note, however that head roll could provide a vertical disparity depth cue (6).

Despite these factors, our subjective experience of apparent depth is unchanged during head movements and when rolling the head towards one’s shoulder the world does not look tilted. Is this sense of stereoscopic depth ‘constancy’ across changes in head and eye position accompanied by constancy in stereoscopic sensitivty? One study investigated the effect of ocular torsions and thus changes of the image orientation relative to the rolling direction (7). These authors created stereograms that use rotational instead of horizontal image offsets to create a stereoscopic depth cue with broadband, binary black-white noise stimuli. Ocular torsions were induced by asking observers to adopt various up- or down gaze directions. While testing different gaze directions in a habitual head position, participants were not able to reliably report depth when ocular torsion directions did not match the stereogram rotation, ostensibly demonstrating that the ability to perceive stereoscopic depth is confined to horizontally fixed regions of the retinae.

To our knowledge, only two studies have examined the effect of head tilt - specifically roll - on stereo acuity (Lam, Cheng, Kirschen, & Laby, 2008; Walchli & Ebenholtz, 1965). Walchli & Ebenholtz (1965) measured depth discrimination thresholds and found a reduction of stereoacuities when increasing the headtilt ‘roll’ from primary position to 90°, a relationship that was well fit with a cosine function. However, that study employed a pair of physical lines within a rotating cylinder and thus horizontal disparity is present but is confounded with line orientation, which has been shown to affect stereoacuity even without head tilt (10). Lam et al. (2008) reasoned that the IPD is directly related to stereo acuity threshold (Schor & Flom, 1969). Using spatially broad-band random dot stimuli and rolling the head up to 30°, they found for approximately 77% of the participants a decrease of stereoacuity.

The current study’s overarching aim was to investigate how the spatial structure of different stereo acuity tests and changes of pitch, roll and yaw head tilts and roll stimulus tilts affect the stereoacuity of binocularly normal-sighted adults.

## 2. Experiment 1: Head Tilt and Sparse-Pattern Stereoacuity

### 2.1 Methods

Written and verbal information about the project were provided in advance to the participants and they gave written informed consent before taking part. Northeastern university’s ethics board approved to conduct the experiments on human participants as the experiment deemed to be in line with the ethical principles of the Helsinki declaration of 1975.

#### 2.1.1 Participants

Ten adult participants with normal or corrected-to-normal vision and stereoacuity of 60”(Titmus) or better were recruited from colleagues laboratory as well as from the undergraduate student population at Northeastern University, Boston. Undergraduates received course credit towards the completion of their Introductory Psychology course in exchange for their participation.

#### 2.1.2 Equipment

##### Head position control

Head position was monitored with a Bluetooth 4.0 Inertial Measurement Unit (IMU; 3-Space Sensor™, Yostlabs, Portsmouth, Ohio, USA) sensor, with resolution<0.08° and 100Hz sampling frequency, fastened to the back of the participant’s head on a GoPro (USA) head strap. The IMU was calibrated and tared with a 2D spirit level. The IMU provided continuous head orientation output as quaternion angles and Euler angles relative to the calibration and taring orientations. Data were sent wirelessly to the computer where a customized C++ program ran simultaneously with MATLAB 2019a using Psychtoolbox-3 and Open-GL to calculate the head location and orientation in real time. Feedback concerning 3D head orientation was presented in real-time to the observer on the experimental display via a 3D 6° colored cube at the center of the screen controlled with Open GL software and by a 2° white head-posture-contingent ring that moved with the observer’s head. The target head posture was indicated with a wire frame cube and by a 6° green ring. To obtain the required head posture, the observer’s task was to move their head to align the cube with the wire frame (at which point the cube changed from red to green), and simultaneously moved the white ring so that it was within the green circle (see Figure 3). Observers were required to hold the required head posture withn ± 3° of the required head posture and fixate the central of the screen (see Eye Movement Control section) for at least 800msec, then an experimental trial was initiated.

**Figure 1:**
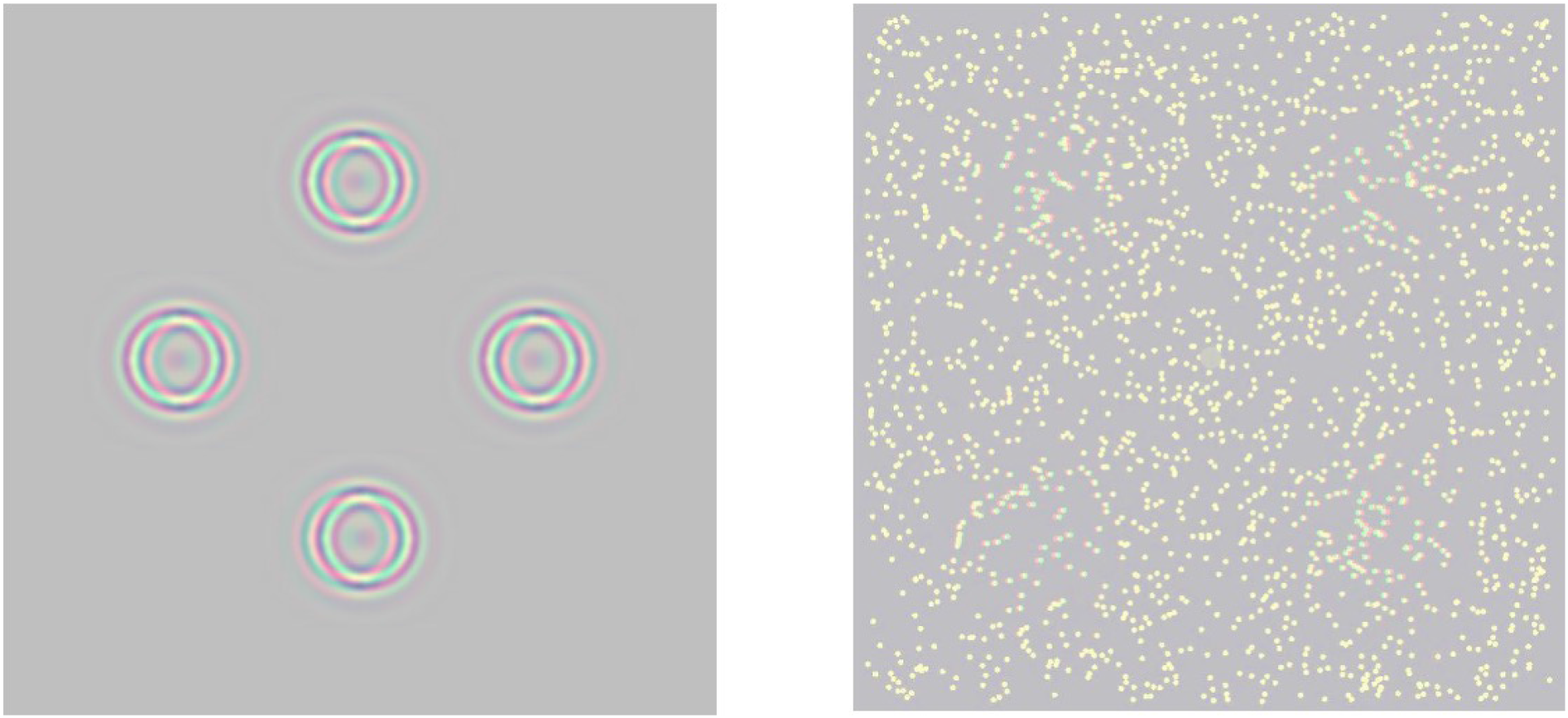
Red-green anaglyphic versions of the stereotest used in the first (left) and second (right) experiment – in the experiment, stimuli were greyscale and stereo was controlled with a polarised display and glasses. A) Circles subtended 1.25° diameter and were digitally bandpass filtered with a log cosine with a peak spatial frequency of 4c/° and Full-Width-Half Height bandwidth of 1 octave. Stereoscopic disparity was generated by horizontal dispalcement of the circles in the left and right eyes, that was under the control of a computer staircase. Three of the circles were in uncrossed disparity, the target circle was crossed disparity, the observer’s 4AFC task was to indicate the location of the target circle by pressing a corresponding computer key. B) Random dot stereograms were created with white Gaussian dots (σ =0.04°) with a density of 30 dots/^°-2^. Stereoscopic disparity was created with a Gaussian (σ=0.67°) displacement of the dots in each eye, that was concave in three and convex in the target quadrant. The observer’s 4AFC task was to indicate the location of the target by pressing a corresponding computer key.

**Figure 2:**
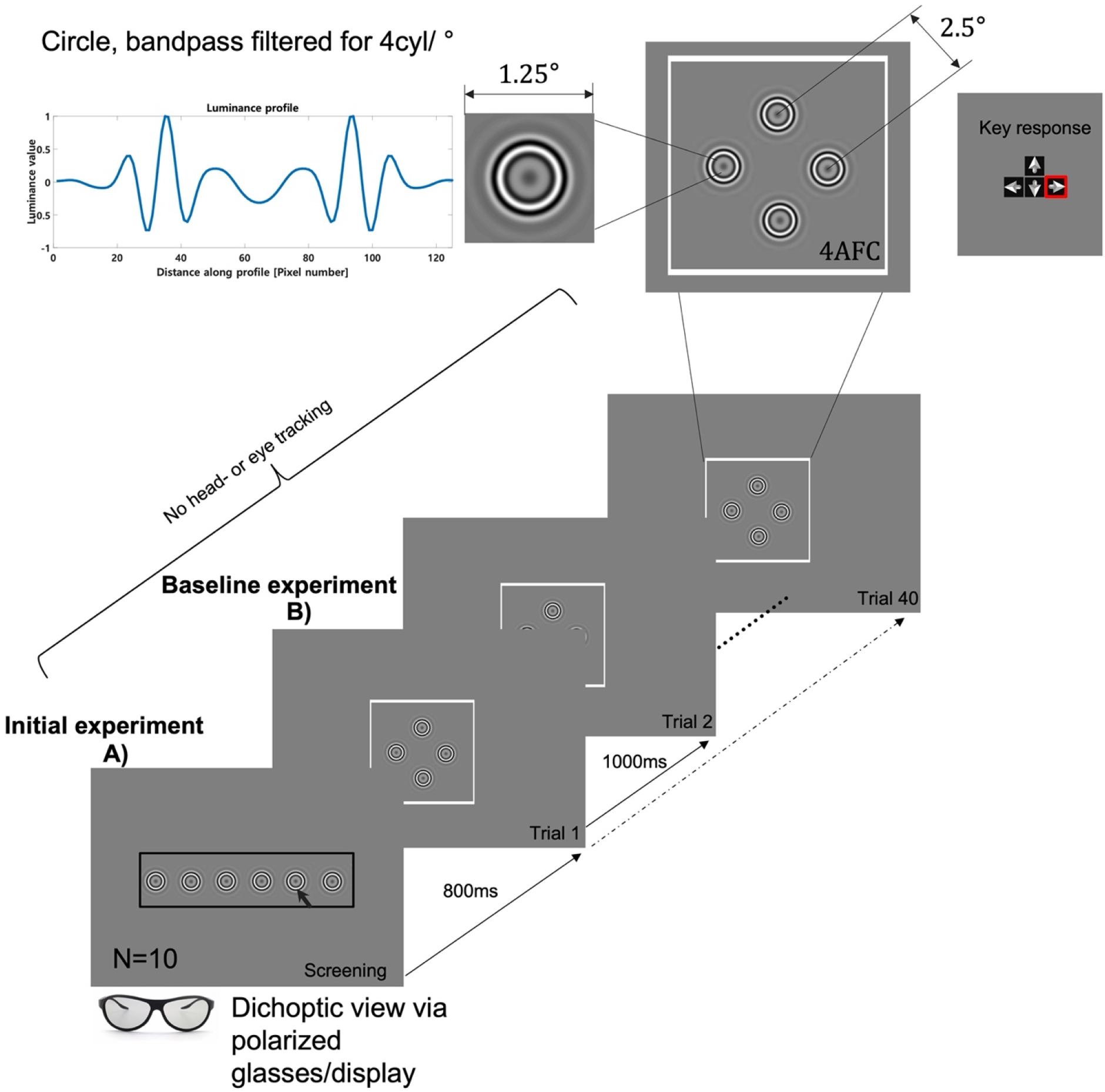
Illustration of the display for stereoacuity measurement with band-pass filtered circle stimuli. A) An initial stereoacuity measure was taken as an estimate threshold in upright, habitual head position. The stereo disparity of 6 bandpass filtered circles decreased from 1500 to 40 arcmin in 6 evenly spaced log steps. Observers used a mouse to click on the rightmost circle that still appeared to be in front of the display. This disparity was used to start the staircase for the forced choice task. B). Four band-pass filtered circles (three uncrossed, one crossed disparity) were presented for 1 sec in each of 40 trials in total for each viewing condition. In a 4 alternative forced choice task with visual (fixation color) and auditory feedback, the participant indicated the circle that appeared in front of the display via keypresses. The graph on the top left depicts the luminance profile of one of the stimuli.

**Figure 3:**
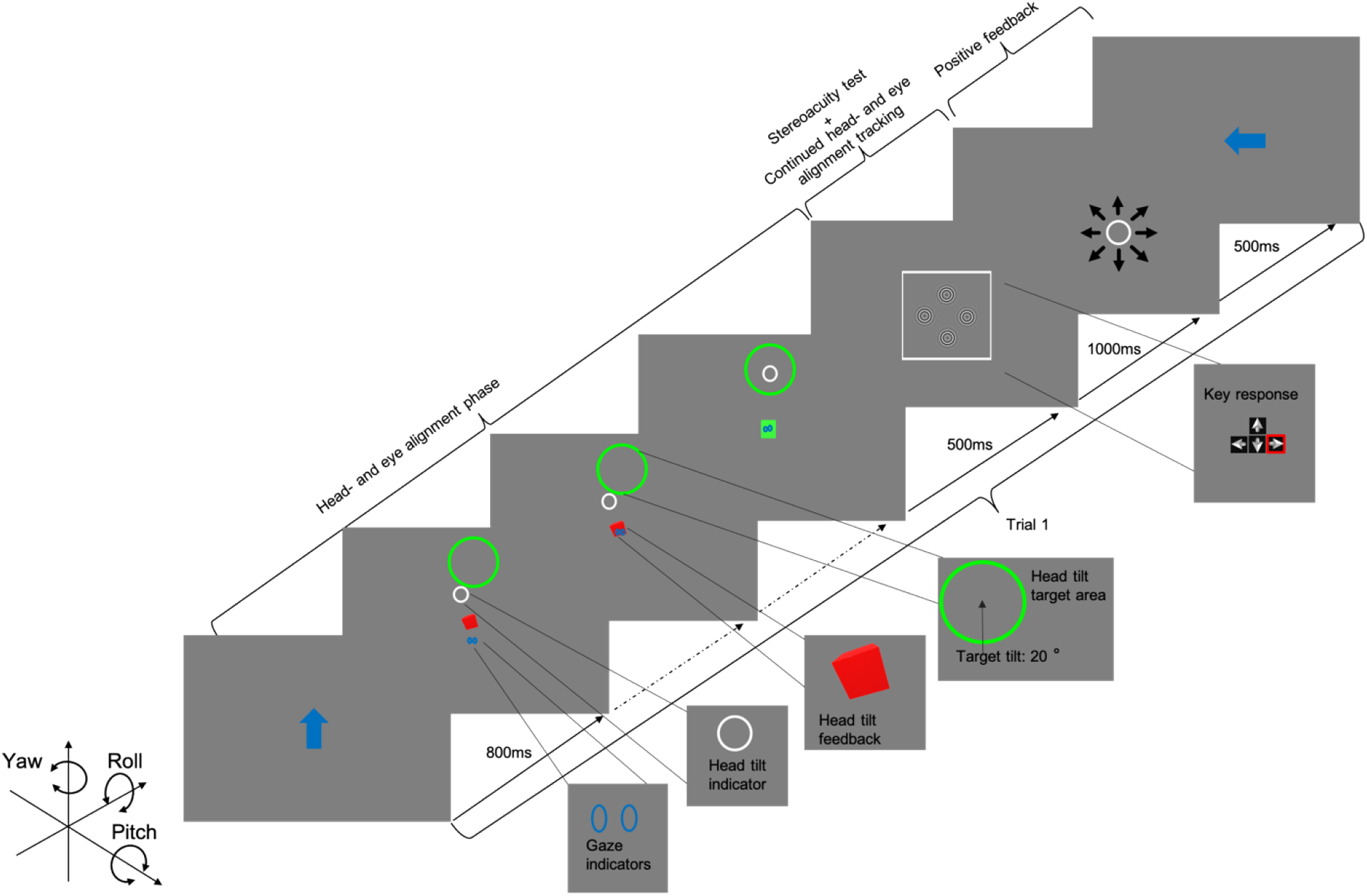
Schematic Illustration of the head and gaze posture paradigm for the first experiment. A trial began with a blue circle that indicated the required head tilt orientation for 800ms. The subject was required to fixate (blue ellipsoids, one for each eye, provided real-time feedback for this task) the center of the cube while tilting their head to move a head-posture-contingent small white circle within the target area (green circle). Once this was accomplished, the cube changed its color from red to green and the subject was required to hold these eye and head postures for 800ms to initiate the stereoacuity task. The stereoacuity screen lasted for 1000ms after which the participant indicated which of the circles was in front depth. During the stereo test, the participant was required to maintain the target eye and head postures; in case eyes, head or both were out of range the trial was aborted and the head and eye alignment phase was repeated for a newly randomised trial.

##### Eye movement control

Binocular eye movements were measured simultaneously with Tobii EyeX (Sweden) remote eye tracker with a sampling rate of 60 Hz, using the Tobii EyeX Matlab toolkit (12). The observer’s gaze task was always to look directly at a fixation point at the center of the screen, regardless of the head posture required for any condition. Real-time feedback concerning gaze direction for each eye was provided with two small circles, one for each eye. The observer’s task was to maintain the gaze-contingent rings within 3° of the fixation target (see Figure 3). Once observers held this gaze and the required head posture for at least 800msec, an experimental trial was initiated.

##### Monitor and computer

Stimuli were presented on a polarized 42” 3D TV from LG (model 42LM6200-UE) with refresh rate of 60 Hz at a resolution of 1920 by 1080 pixels. Dichoptic viewing was obtained with LG cinema 3D glasses (AG-F310), over spectacles if worn. Crosstalk was minimized with Psychtoolbox’s *StereoCrosstalkReduction* and *SubtractOther* routines to minimize the subjective visibility of a 100% contrast 4 c/deg sine grating presented to one eye and a mean luminance field presented to the other eye. The monitor was gamma-corrected prior to the experiment using a Photo Research SpectraScan 655 (Norway) spectrophotometer. The viewing distance was set to 120 cm from left eye’s corneal apex to the screen during the upright baseline condition.

#### 2.1.3 Stimuli

The stimuli were generated using Matlab (2019a) with Psychophysics Toolbox (13) extension on a Microsoft Windows (Version 10.0.18362.1). Stimuli were presented on a grey background with a mean luminance of 25 cd/m^2^. The pixel size was 0.734mm^2^.

##### Stereoacuity stimuli

#### 2.1.4 Procedure

##### Stereoacuity Screening

There are large individual differences in stereoacuity, even in the typically sighted population (14). We therefore implemented an initial screening step (15), to individualize the starting parameters of the staircase for each observer. Six bandpass-filtered circles were horizontally positioned with a center-to-center distance of 2.5° (Figure 2A). The diameter of each circle was 1.25° and was filtered with a log cosine filter with a peak spatial frequency of 4cyl/° and Full Width at Half-Height bandwidth of 1 octave. The horizontal disparity of each circle decreased from 1500” to 40” in six logarithmic steps from left to right. Observers were required to click on the rightmost circle that still appeared to be in front of the display. The disparity of this circle was used as the starting value of the staircase in the main experiment.

Next, stereoacuity was measured with bandpass-filtered circles as for the initial trial except: four circles were arranged in a diamond shape and presented centrally at a primary gaze and head position, with 2.5° distance between each circle’s center (Figure 2B).

##### Stereoacuity Measurement

Following stereo screening, stereoacuity was measured with a four alternative forced choice (4AFC) procedure. Four bandpass-filtered circles (peak spatial frequency 4cyl/°) arranged in a diamond-configuration were presented for 40 trials (Figure 2). Three of the circles were in uncrossed disparity (i.e behind the surface of the display), the target circle in a random position each trial, was in crossed disparity (i.e. in front of the screen). The size of the disparity on the first trial was equal to the observer’s screening threshold, and was under the control of a staircase procedure (16) on subsequent trials. Disparity was decreased by of 1/3dB following a correct response or increased by 1dB following an incorrect response. Once observers had attained the required head posture and their gaze was at the center of the screen, the 4AFC stimuli were presented for 1000msec. The observer’s task was to indicate the location of the target (the circle with crossed diparity) by proessing a corresponding key, while maintaining head and eye posture. Audio and visual feedback (high pitch beep and green fixation point following correct responses or low pitch beep and red fixation point following incorrect responses) were provided to aid participants to maintain their attention and motivation for the task.

Stereo acuity was measured under 7 conditions: *BaselineStereoacuity* in which observers adopted natural head and eye posture. The stereoacuity threshold from the baseline condition was used to initialize the staircase for measuring *Head-tilt Stereoacuity:* in which observers were required to adopt 6 head eye postures: ±20° pitch, roll, or yaw in random order each trial of 40 trials. An area tolerance for central fixation and that of head tils were ± 3°. For example, during the Up task, pitch angles could vary from −17° to −23°, while yaw and roll angles could vary from −3° to +3°.

#### 2.1.5 Data analysis

All graphical representations of results were created with Matlab 2021a. Specific details concerning each analysis are provided below.

##### Stereoacuity

The preliminary stereoacuity threshold determined with the initial trial was computed in the following way using a logistic function:

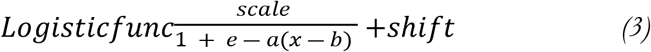

The disparity (scale) that a participant choose was converted to log_10_ arcsec disparity, the scale is defined as 1 – 6AFC^−1^, the shift is defined as 6AFC^−1^. The parameters *a* and *b* are obtained as estimates of data making use of the Matlab *fminsearchbnd*, which searches for a minimum problem specified by

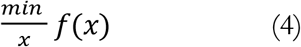

in which *f*(*x*) is a function that returns a scalar (*a*), and *x* is a vector or a matrix (*b*), with lower and upper bounds constraints.

A cumulative Gaussian was fit to the raw data of proportion correct as a function of stereoscopic disparity from which stereoacuity threshold was estimated at the 62.5% correct point *p*:

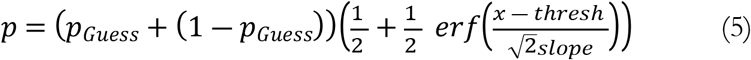

where *p_Guess_* was 25% (4AFC task), *x* is disparity, and threshold and slope were free fit parameters. Stereothresholds, here refered to as stereoacuities for 7 headtilt conditions were calculated for each of the ten participants and were stored in an 7*10 matlab array. One-way Analysis of Variance (ANOVA) using Matlab 2021a was deployed to compared results for an effect of headtilt condition.

##### Headtilt

Each observer’s angular data were averaged across trials and in addition we calculated the mean and medians across observers for the duration of the experiment (1000ms) and depicted the data in a line plot.

##### Eye tracking

The vertical and horizontal data were extracted for each eye, condition, trial, were then averaged across trials and observers, and finally the results for each condition are depicted in a line subplot. The mean data for each oberserver were used to calculate the standard error of means and were depicted using the Matlab function shadedErrorBar (17). We calculated the angular eye location biases as averages across trials and observers for each condition’s left and right eye location biases and used one sample t-tests between eyes.

### 3. Results

#### 3.1 Head location

Figure 4 shows the head posture for each condition, up or down pitch, left or right yaw and clockwise or counterclockwise roll averaged across trials, including each observer’s single data and an average across observer (thick red lines). Data for each participant are shown by line color, the solid blue line indicates the mean across observers. As required by the paradigm, observers were able to attain and hold a head posture of ±20° pitch, roll, or yaw throughout the test interval, with a median head tilt of 19.7° calculated across trials, participants, and conditions.

**Figure 4:**
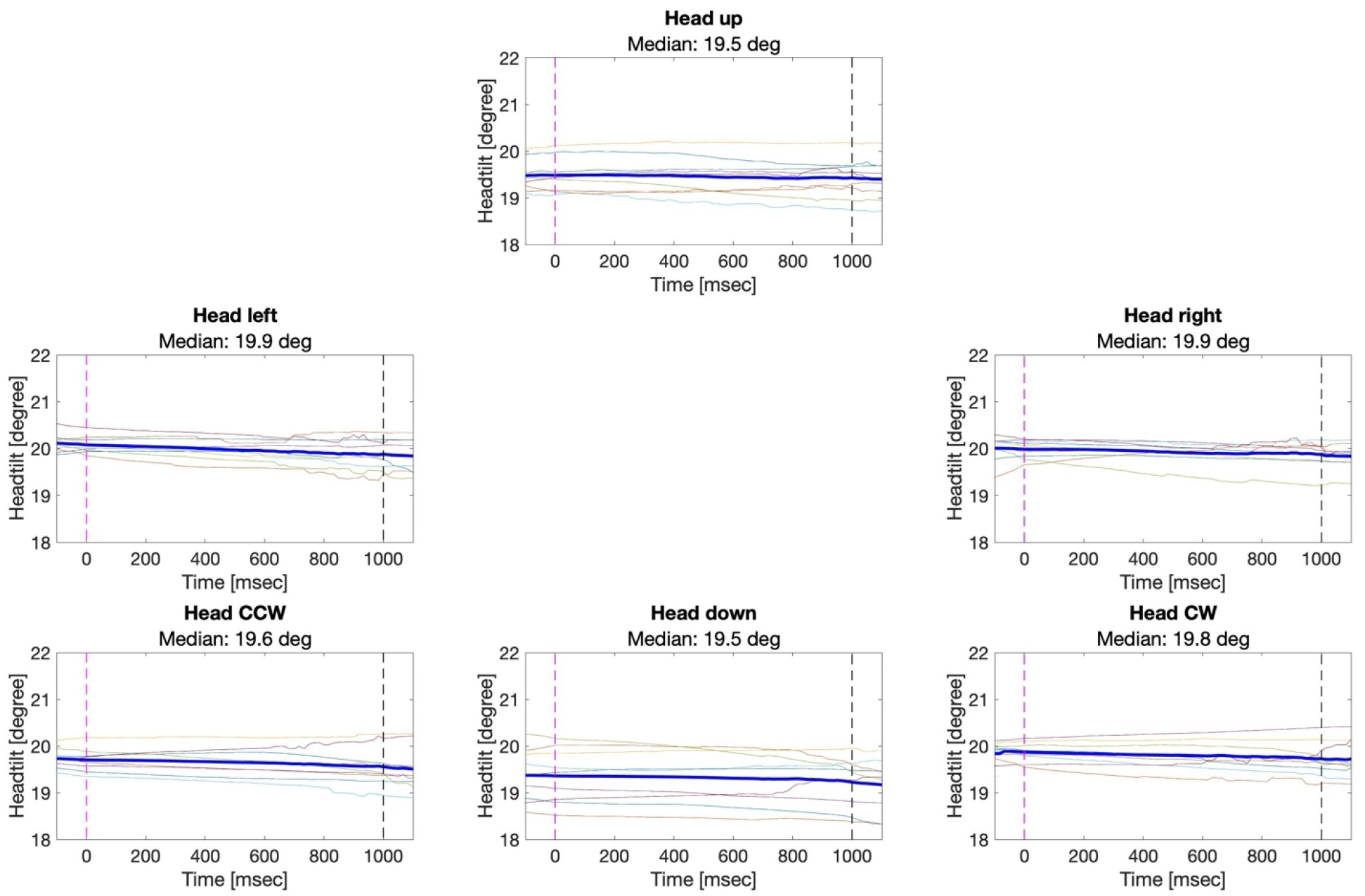
Head tilts (ordinate) plotted separately for each head posture condition across time (abscissa). The vertical dahsed lines at Time 0 and 1000msec indicate the start and end of the 4AFC stereoacuity task, respectively. CCW and CW refer to counterclockwise and clockwise head tilts, respectively. The differently colored thin lines depict each participant’s mean across trials, the means across participants are indicated using thick blue lines. Median head tilt angles for the trial period (0 to 1000ms) are shown in the subtitles.

#### 3.2 Changes of horizontal and vertical eye movements during head tilt

Figure 5 shows the estimated horizontal direction of gaze of the right (red) and left (blue) eyes for each head tilt condition (up or down pitch, left or right yaw and clockwise or counterclockwise roll) averaged across trials and observers, shaded areas indicate ±1 standard error of means. The overall bias towards left or right off the center of the screen was with −1.76°±1.01 for the right and 1.88°±0.94 for the left eye not statistically different t(5)=0.22, p=0.84. Although observers maintained binocular gaze on the center of the display where the stimulus was presented, changes in head posture caused a decrease in the planar interocular distance relative to primary gaze. For pitch (head down), these are expected convergence eye movements that are on average 2.0 °±0.6° SD, consistent with previous reports (18).

**Figure 5:**
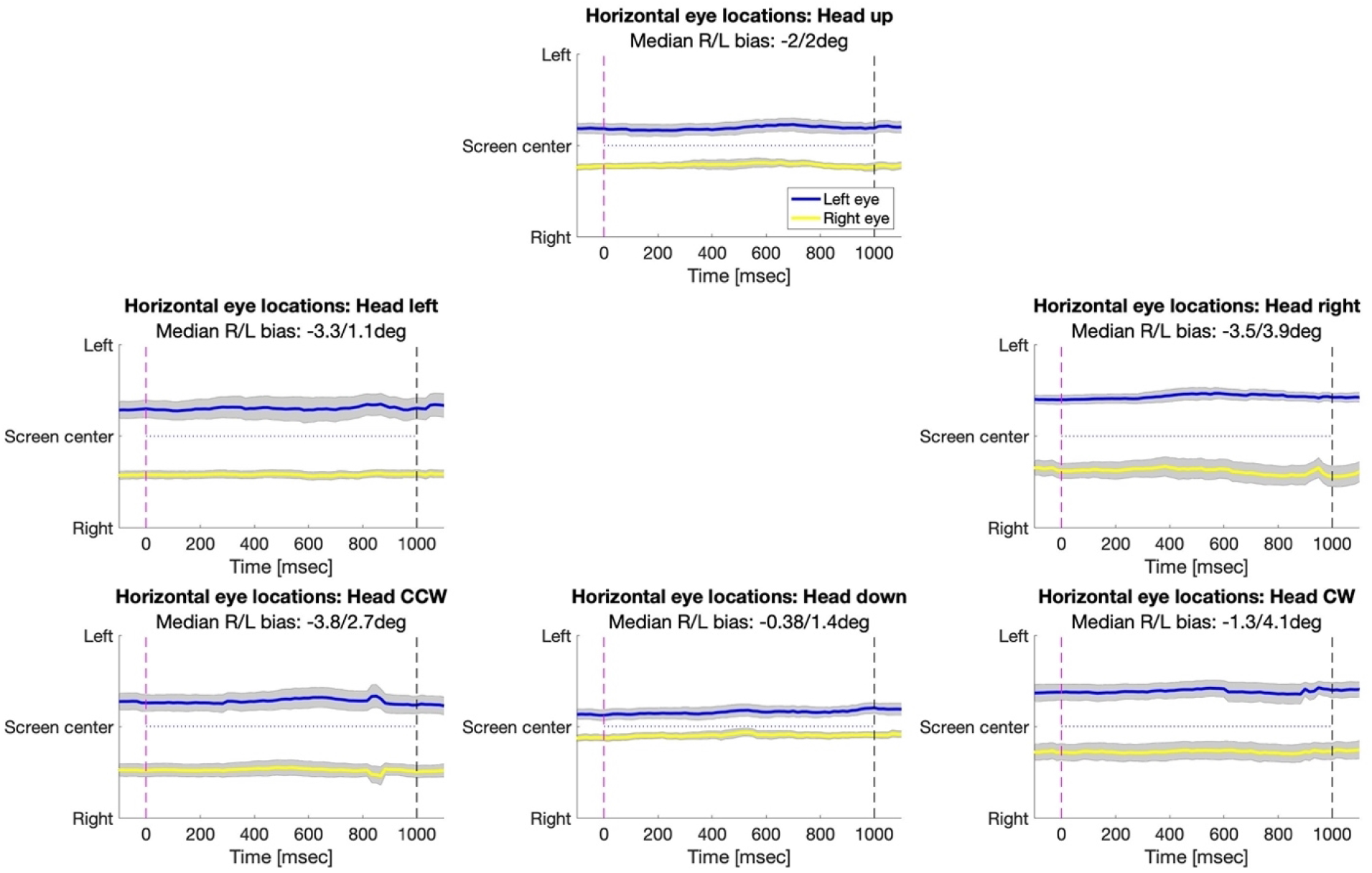
Eye tracker estimates of the horizontal gaze direction (ordinate) relative to the screen center (0°) of the right (yellow) and left (blue) eyes plotted separately for each head posture condition across time (abscissa). Grey shaded regions indicate the respective standard deviations. Positive gaze direction values indicate the right screen half, negative value indicate the left screen half in degrees of visual angle. The vertical dahsed lines at Time 0 and 1000msec indicate the start and end of the 4AFC stereoacuity task, respectively. The median biases for the duration of the experiment for the left (blue, top horizontal line) and right eye (yellow, top horizontal line) and their standard error (grayshaded thick line) are shown in the captions. Although observers maintained gaze on the center of the display, changes in head posture caused a decrease in the interocular distance (leftward movement of the right eye and rightward movement of the left eye).

Figure 6 shows the estimated vertical direction of gaze of the right (red) and left (blue) eyes for each head tilt condition (up or down pitch, left or right yaw and clockwise or counterclockwise roll) averaged across trials and observers, shaded areas indicate ±1 standard error of means. The overall bias towards upper or lower half of the center of the screen was with 1.47°±0.96 for the right and 1.42°±0.87 for the left eye not statistically different t(5)=0.12, p=0.91. As expected, changes in head posture casued a change in the planar estimate of vertical gaze for roll while there was no change in vertical position for pitch up and down as well as yaw left or yaw right. For counterclockwise roll, the estimated direction of gaze for the right eye shifts up and the left eye shifts down, with the opposite occurring for clockwise roll.

**Figure 6:**
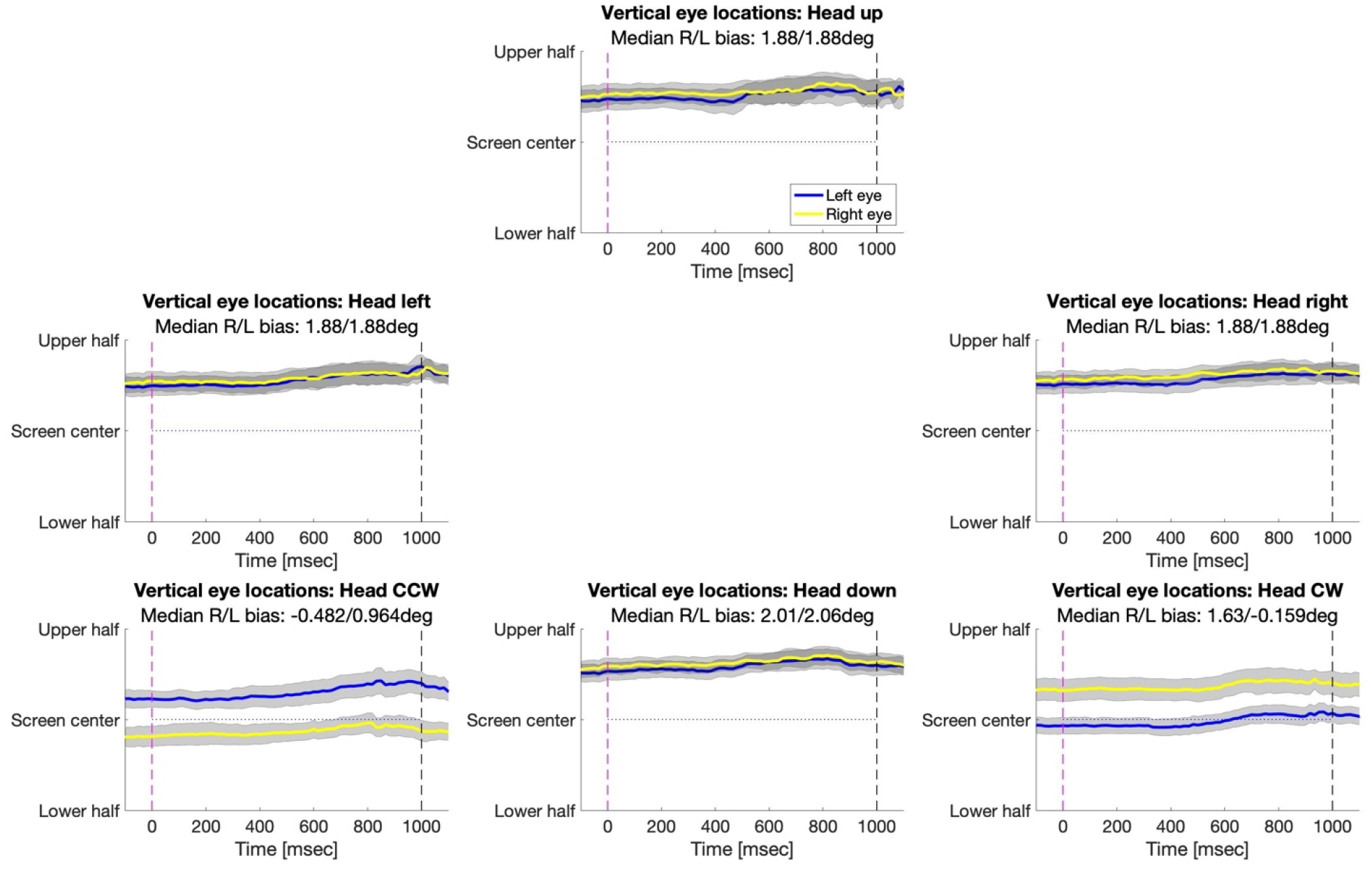
Eye tracker estimates of the vertical gaze direction (ordinate) relative to the screen center (0°) of the right (yellow) and left (blue) eyes plotted separately for each head posture condition across time (abscissa). Data are plotted as in Figure 5, except for vertical position.

#### 3.3 Effect of head tilt on stereoacuity

We compared the stereoacuity outcomes for the different head tilts (Figure 7) using a repeated measures one-way ANOVA and found no statistical significant difference between the various head tilts conditions [F(6,63) = 0.12, p=0.994]. Even though participants attained the required head tilt (median= 19.7°) and there was a predictable change in horizontal and vertical interocular distance. The mean stereoacuity, across subjects and conditions was 122 arcsec ±3.5 standard deviation and did not significantly vary with head posture (see Figure 7). This estimate is in line with typical estimates of stereo acuity for normally-sighted observers with good stereoacuity, who have a peak at 96 arcsec (14). The lack of effect of head tilt on stereoacuity is remarkable given that head tilt decreases interocular distance and is most surprising for head roll, which caused a decrease in horizontal interocular distance and a deviation in vertical alignment, both of which should cause a decrease in stereoacuity.

**Figure 7:**
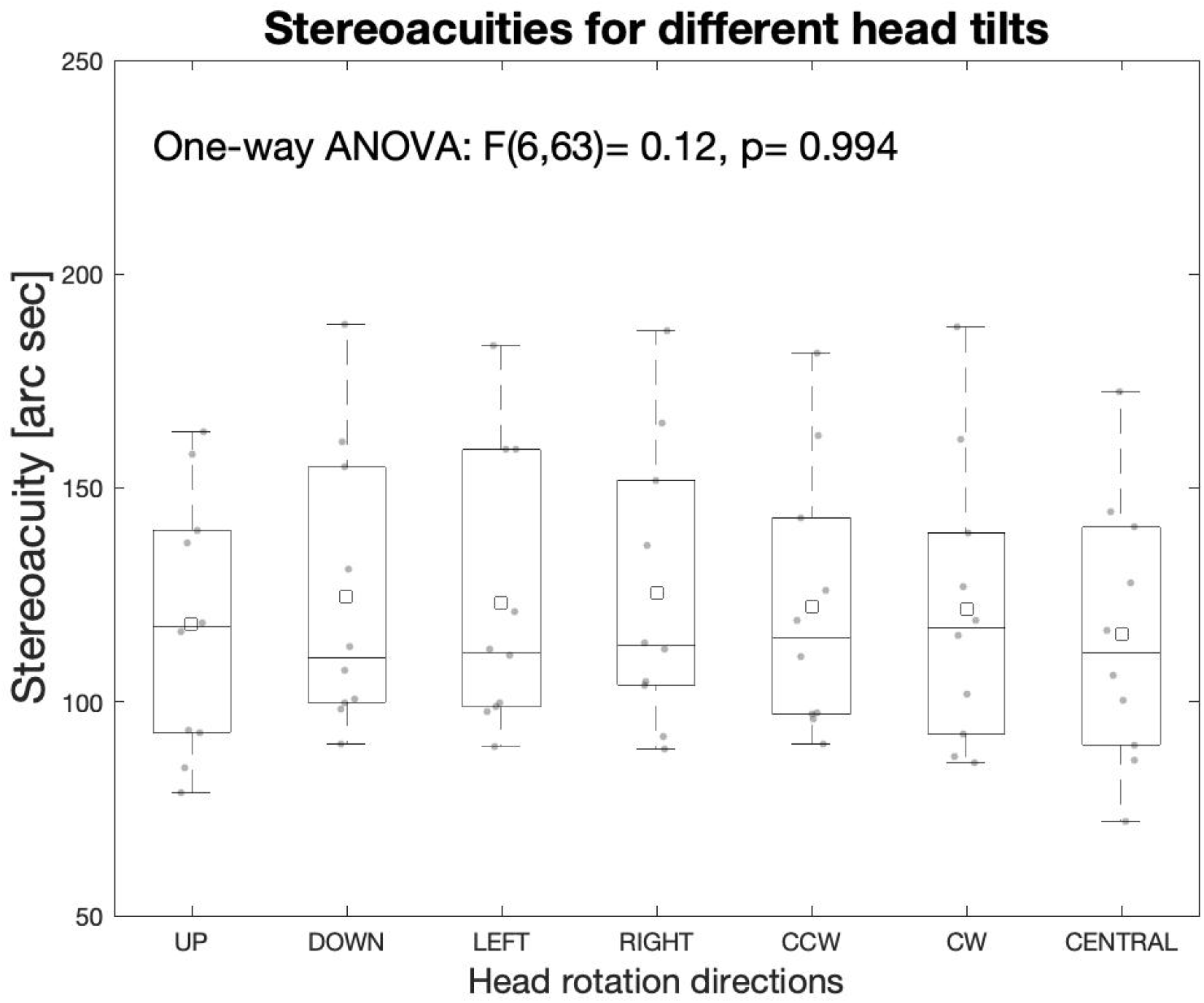
Stereoacuity threshold for different head tilts. Stereoacuity thresholds in degrees of arc are depicted on the y-axis shows while the x-axis shows head tilt orientations including pitch up and down, yaw left and right, roll clock- and anti-clockwise, and central with thresholds from the baseline experiment. The black squares depict the means, the red lines the medians, the blue boxes the 25% to 75% interquartile ranges, and the whiskers the most extreme threshold values. Each blue data point represents individual’s mean, averaged across trials. The one way ANOVA result is shown on top.

**Figure 8:**
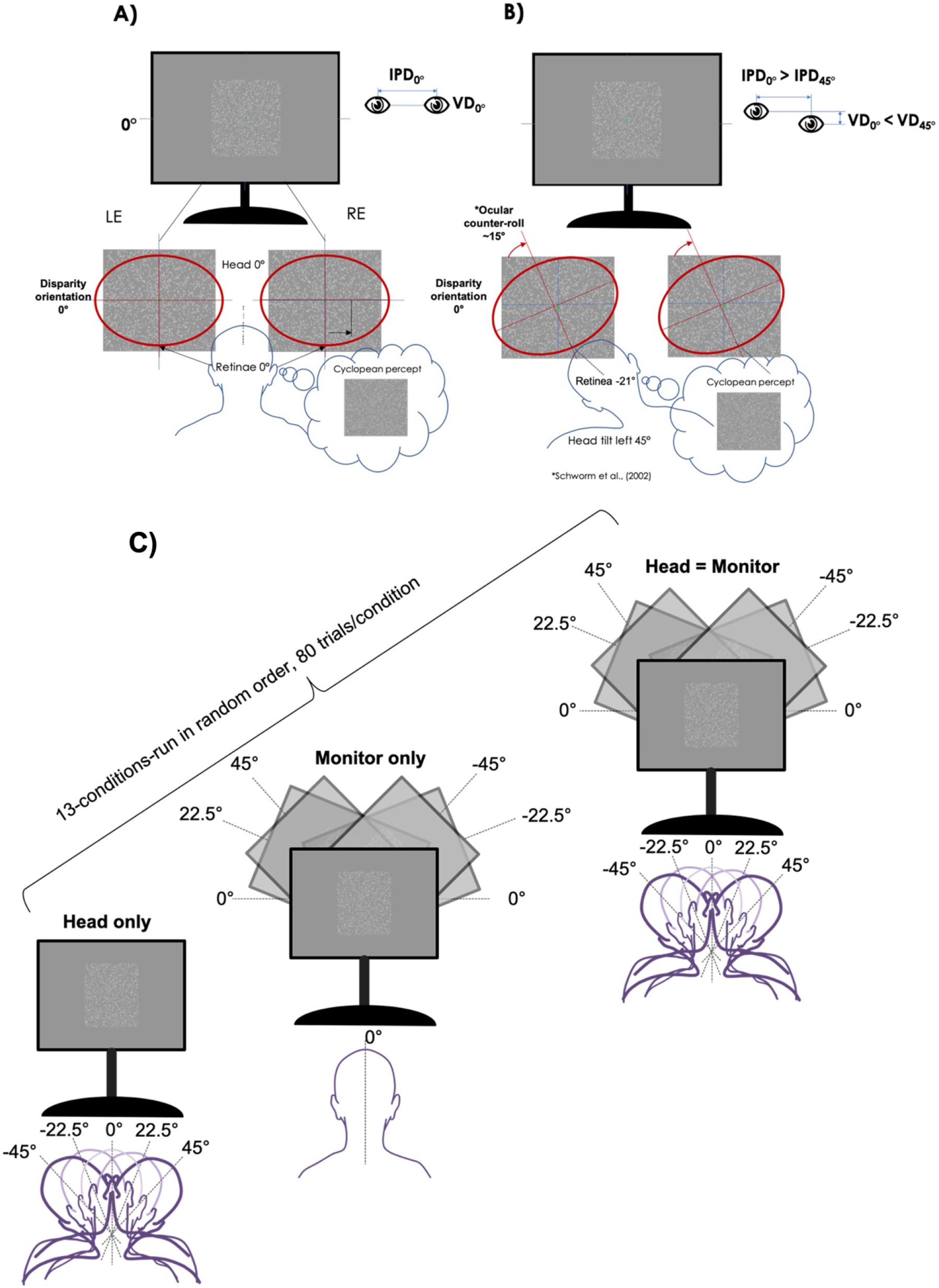
Illustration of the interaction between head roll and its influence on retinal image representation. A) Upright head position in primary gaze for a random-dot stereo test. B) 45° left head tilt causes a 29% reduction of the IPD and increased vertical distance (VD) between left and right eye, and an ocular-counter roll that compensates only approximately 15° of that of the head tilt (21). C) Schematic of the procedure used in the 2nd experiment. Participants completed 80 trials per 13 different head tilt positions in randomized order. Breaks were encouraged between trials. Participants were instructed to reach the tilt position by only tilting the head via neck while keeping the torso upright.

## 4. Experiment 2-Head Tilt and Random Dot Stereoacuity

Yaw and roll reduced the IPD but roll also alters the horizontal correspondence centers of the eyes due to deconjugated OCR (19), thus both would be expected to reduce stereoacuity. In contrast to these predictions, in Experiment 1, we found that stereoacuity did not significantly change when titling the head 20 degree using narrowband circle stimuli.

Lam *et al*. (2008) reported that about ¾ of their participants experienced a decline in stereoacuity when rolling the head, but about ¼ did not and our results are in agreement with the latter subset of these subjects. Lam et al employed the circle stimuli of the Binocular Video Acuity Tester, similar to Titmus circles and the stimuli employed in Experiment 1. Such stimuli contain sparse elements and have a relatively narrow spatial frequency spectrum. Other stereoacuity tests employ random dot stimuli, which have a broader amplitude ‘white’ spectrum that is dominated by information and high spatial frequencies. We have previously demonstrated that high spatial frequency energy in random dot stimuli can mask information at other spatial scales (20). Furthermore, for the detection of binocular disparity, dense random dot stimuli contain potential false correspondences along the horizontal axis that are not present in the sparse circle stimuli. In Experiment 2, we repeat the main conditions of Experiment 1 in order to examine whether stereoacuity remains invariant of head posture for dense random dot patterns.

### 4.1 Methods

#### 4.1.1 Participants

Eight adult participants (3 males, 5 females) with an average age of 29 ± 3.4 years carried out the experiment. All participants except author J.S. were naïve to the purpose of the study. All other aspects were as for the first experiment.

#### 4.1.2 Head and monitor position control

To measure the head and monitor tilt when rolling to the left or right, we used a protractor with a combined spirit level placed at the center of the polarizing glasses (for head tilt) or at center of the upper monitor frame (for monitor tilt), respectively. We then tilted head and monitor accordingly to the desired angle of tilt. We did not record eye movements for this experiment, since the observer’s task was always to look at the center of the screen where the stimuli were presented.

#### 4.1.3 Stimuli

The stimulus size was 8°*8°, made of four 4°*4° quadrants, each containing white (50 cd/m^2^ peak through the polarised glasses) Gaussian dots (σ=0.04°), with a density of 30 dots/°^−2^ presented on a 25 cd/m^2^(through the polarised glasses) grey background (Figure 9). Interocular horizontal disparity was created by shifting dots in the left and right eyes in opposite directions according to a Gaussian (σ=0.67°) whose peak amplitude was controlled by a staircase (16). Offsets were crossed/convex in three and uncrossed/concave in one quadrant and were aligned with the azis of the monitor (horizontal for 0° monitor tilt). The fixation point in the center was a 0.25° by 0.25° square.

**Figure 9:**
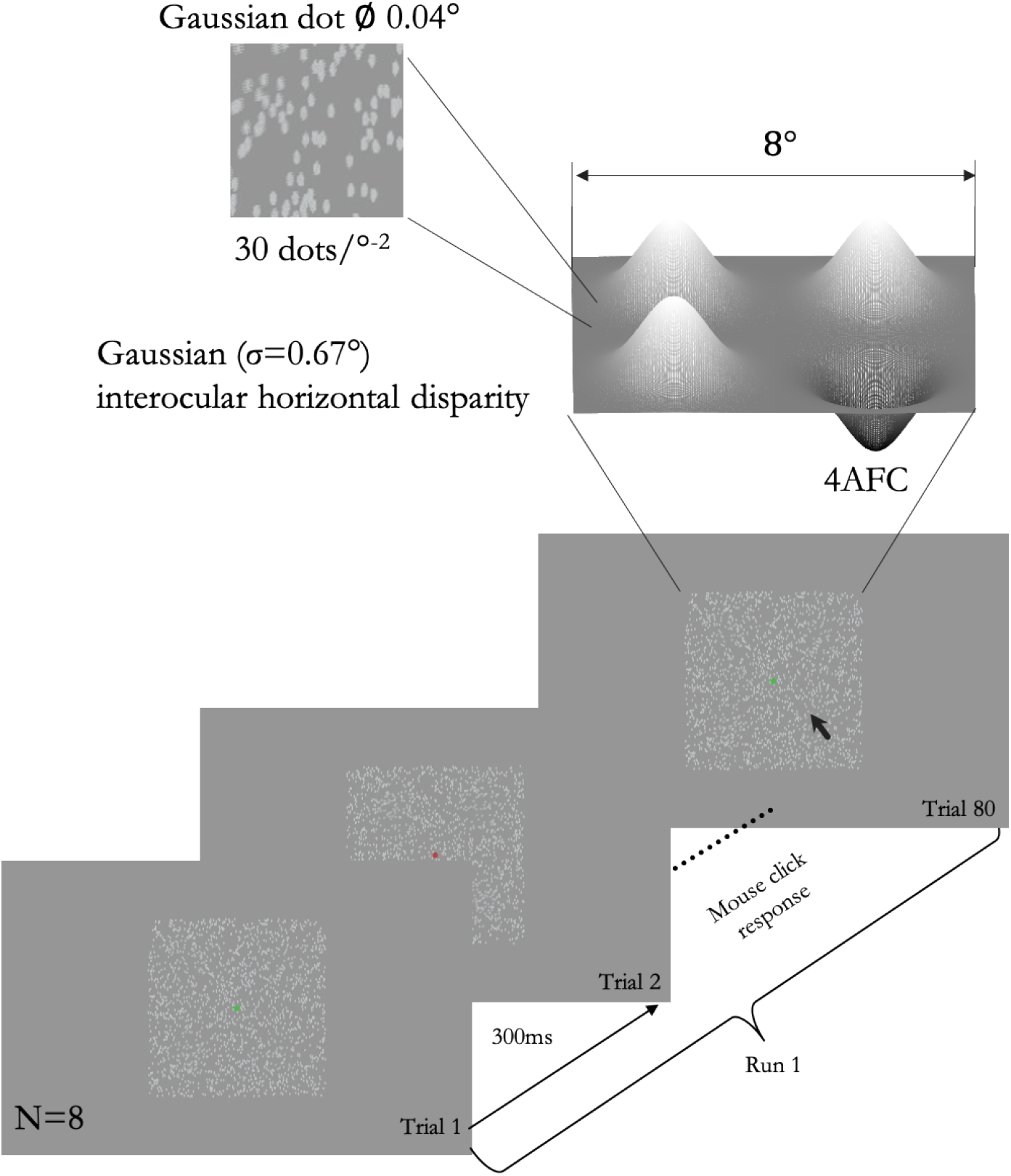
Illustration of stimuli and task for Experiment 2. The locations of random gaussian elements (σ=0.04°) were adjusted in each eye to create four Gaussian (σ=0.67°) depth profiles, three with convex/crossed disparityand one with concave/uncrosed disparity. Each run consisted of 80 trials, each lasting 300ms. The observer’s 4AFC task was to click on the concave region, feedback was provided by a green fixation spot and a high pitch beep for correct reponses, or a red spot and low-pitch beep for incorrect responses.

#### 4.1.4 Data analysis

We fitted a cumulative gaussian psychometric function to the proportion of correct responses at each disparity for each participant and tilt condition, stereoacuity was estimated at the 62.5%-correct point. We calculated the averages across trials and observers and then fitted the values with cosine functions for each condition using Matlab’s *fittype* and *fit*,

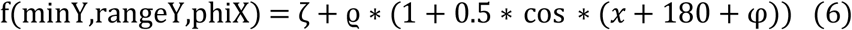

where *ζ* is peak stereoacuity, ϱ is stereoacuity range, *x* represents the headtilt orientation, and φ is the head tilt with peak stereoacuity.

We applied a mixed model ANOVA to investigate the effect of tilt angle, tilt condition, and observers using Matlab’s *anovan* function. Planned multi-comparisons between the conditions were calculated with the *multcomp* function.

#### 4.1.5 Procedure

Participants were seated in a comfortable chair and placed their head into a chinrest while sitting upright. Head- and monitor tilts were determined by the experimenter before each condition using a protractor with a combined spirit level. Stereoacuity was measured while either the head, the monitor, or both were tilted in 0° (baseline), 22.5° or 45° towards left- or right in random order (see Figure 8C). The 4-AFC task for the participants was to determine via mouse click, which of the four targets appeared concave, guessing if uncertain. Interocular disparity was controlled with a staircase (22) that decreased disparity by of 1/3dB, following a correct response or increased disparity by 1dB following an incorrect response. The experiment included 80 trials per condition, two repeats per condition, for 13 conditions in random order. The presentation duration was 300ms per trial and an overall average duration of 1.13sec ±0.01 standard diviation until a response, resulting in approximately 90min testing duration for each participant.

## 5. Results

### 5.1 Effect of head and stimulus tilt ‘roll’ on stereo acuity

All participants experience a decrease in stereoacuity when tilting head or monitor in comparison to the baseline upright position (32.4arcsec ± 1.8 arcsec). There was a significant interaction between conditions and tilts [F(8, 56) = 4.9, p=0.001], which was due to decreasing stereoacuities with increasing head or monitor title [p<0.05], but not tilt of both [p>0.05]. Except for one subject’s results during the headtilt condition, all stereoacuities worsened when tilting the head or monitor whereas for only 3 out of the 8 participant’s stereoacuity slightly worsened during the combined head and monitor tilt condition. Within observer interactions were found for tilt angle [F(28, 56) = 1.8, p=0.03] and tilt condition [F(14, 56) = 2.8, p=0.004].

Compared with the results during the baseline condition using band-pass filtered ring stimuli from Experiment 1 (32.4arcsec), stereo acuity was significantly worse for random dots 115arcsec planned, two-tailed comparison t(16)=7.1, p<0.001. A comparison of the head roll results of the 1st experiment using narrow-band stimuli at 20° (122arcsec) and the results of the 2nd experiments broad-band random-dot stimuli (44arcsec) at 22.5° using a two-tailed independent t-test also showed a significantly worse stereo acuity for the narrow band circle condition t(34)=8.7, p<0.001. Previous results have found no difference between contour and random dot stimuli (23) for upright head postures. However, stereoacuity for random dot stimuli depends strongly on dot size and density (Read & Cumming, 2019), which could account for the difference we find.

## 6. Discussion

### 6.1 Effect of headtilt on stereoacuity

#### Interpupillary distance, Vertical distance and stereovision

Standard tests of stereoacuity are administered for vertical conditions in primary gaze. However, in the real world and for patients with atypical head (torticolis) or eye (strabismus) posture, depth judgements may be made with the head tilted along at least one axis of pitch, roll and yaw, and with corresponding changes in eye-in-orbit posture. Upward pitch, roll and yaw of the head decrease inter-pupillary distance (IPD), which correspondingly decreases interocular retinal disparity. Head roll also introduces vertical interocular offset, which deviates the axis of horizontal interocular disparity. Note that this axis is further affected by torsional ocular counter roll that partially compensates for head roll, but changes the orientation of binocular receptive fields. In spite of these changes in retinal disparity, we are generally unaware of any change in the apparent depth structure of the environment and we maintain effective hand-eye coordination in 3D. It is possible that this depth constancy is maintained, at least in part, by spatial and dynamic non-stereoscopic cues to apparent depth. In this study, we examined whether stereoacuity is affected by head and eye tilt in the absence of spatial and temporal monocular depth information.

Experiment 1 showed that stereoacuity did not change significantly with head tilt in any axis when measured with sparse narrow-band circle stimuli in 10 visually healthy adults. This result is remarkable, given that head tilt decreases IPD and retinal disparity. Furthermore, head yaw introduces a small interocular difference in distance from the target object and thus changes the focal points of the two eyes as well as retinal image size, but those factors appear to have a negligible impact on stereoacuity in our study.

Several studies (25,26), including a large population study (*n*=*1060*) have found a negative correlation betweeen IPD and stereoacuity, where greater IPDs tended to have worse stereoacuity. In those studies, however, head orientation was primary and the test patterns were based on dense random dot stimuli, whereas in our study heads were tilted and stimuli were sparse. We hypothesized that there could be at least two explanations of the difference between results. Firslty, the visual system could have a mechanism that adaptively changes binocular correspondence fields to compensate for head tilt. Such a mechanism could adapt to the head tilt in our study, but not in previous studies with fixed head and varied IPD. Secondly, without such a mechanism when the head is tilted, random dot patterns introduce false dot correspondences within binocular correspondence fields, whereas sparse circle stimuli do not. Either of these could in principle explain the different effects of IPD reduction on stereoacuity.

We tested these hypotheses in Experiment 2 by studying the effect of head roll with random dot stimuli. The results showed that stereoacuity decreased monotonically with head or monitor roll (Figure 10). This finding is consistent with the simpler second hypothesis that differences between stimuli account for the differences in the effects of IPD on stereoacuity, and suggest that false dot correspondences impair stereoacuity with head tilt due to unmatched horizontal dot patterns. Nevertheless, horizontal disparity is still reduced in head tilt even with our sparse, narrow-band stimuli, yet had no significant effect on stereoacuity (Figure 7).

**Figure 10:**
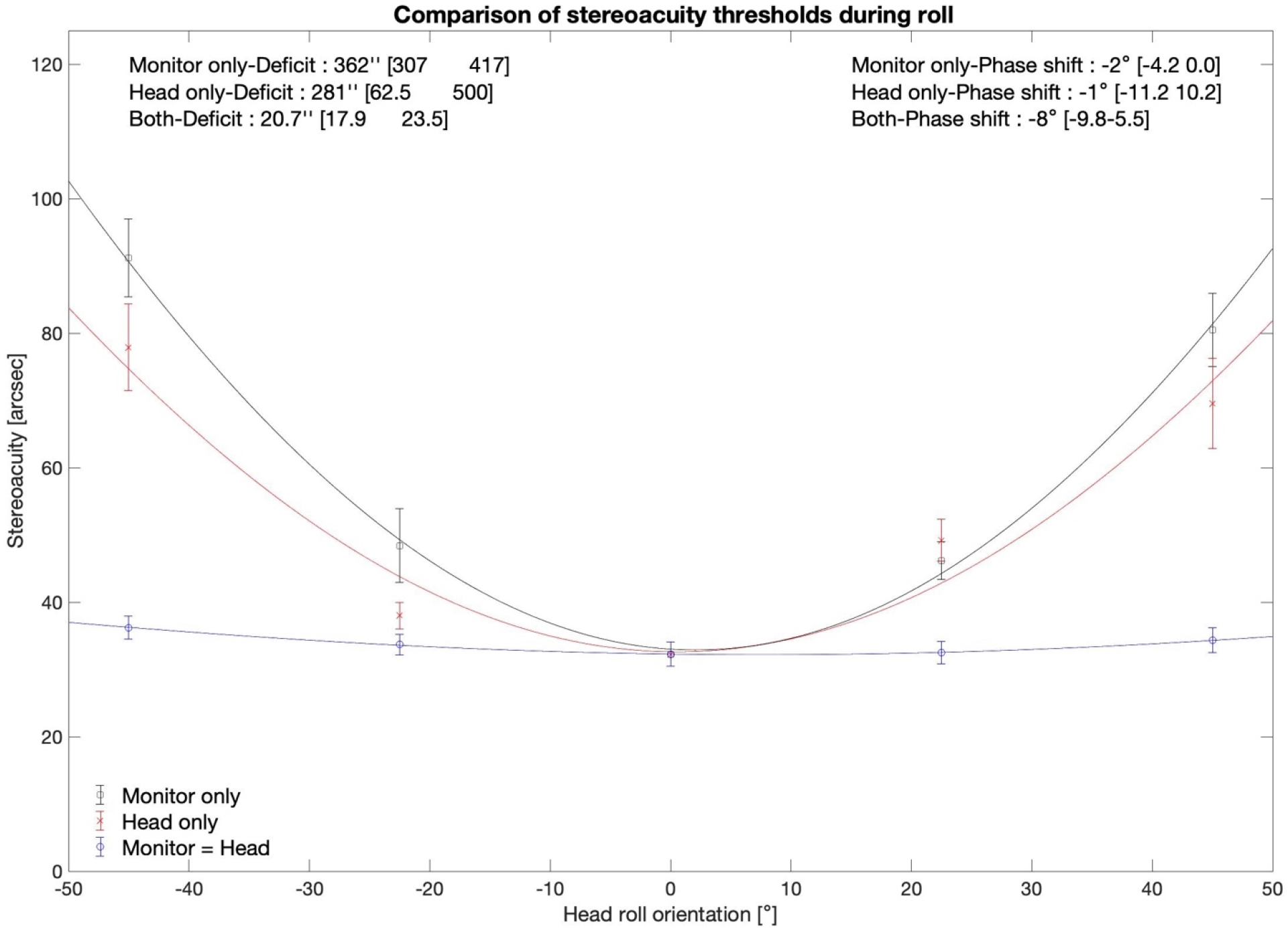
Stereoacuity thresholds as a function of head- and monitor tilt orientation. The x-axis shows the orientation of the monitor (black), head (red) or both (blue), where negative numbers indicate leftward tilts and positive numbers indicate rightward tilts, zero indicated the upright position. Data points represent averages across trials and participants, the error bars represent ±1 SEM, curves show the best fitting cosine functions.

#### Reasons for lack of decline in stereovision with sparse stimuli

In a single-cells recording study using monkeys, Daddaoua and colleagues (2014) measured receptive fields in layer 2 and 3 of area V1 cells before and after tilting the head and found that a subset of neurons shifted their receptive fields on the retina and moreover changed their preferred axis orientation. This finding suggests that there may be an neural mechanism that compensates for head tilt by updating V1 receptive fields, implying that this mechanism involves non-retinal information on eye position to compensate the visual consequences of head and eye tilt. The authors speculate that this mechanism may be involved in ocular counter rolling. The results of Experiment 1 with sparse stimuli are consistent with the existence of a similar mechanim in humans. Standard energy models of disparity coding (28–30) utilise phase or position-shifted receptive fields in which maximum phase disparity and therefore maximum stereo acuity occurs along the horizontal axis. Off-axis phase disparities are also present and can contribute to sterescopic judgments (28,31), but are at lower disparity. It is possible that head tilts interact with sensitivity to these off-axis disparity cues. Given these theoretical considerations and in the light of the results of the current study, Schreiber et al. 2002’s finding that stereovision is eye-fixed in zones of stereo disparity (7) may only be true for broad-band stimuli that contain false correspondences.

#### Future directions

Future studies may investigate the influence of high and low-spatial frequencies on stereovision during head rotations as well as the influence stimulus properties such as the spatial and temporal stimulus structure on ocular counter-rolling characteristics. Monocular occlusion, used as a standard treatment of amblyopia, is known to affect ocular torsions, specifically induced an excyclophoria and vertical phorias in the majority of subjects (32). Indeed, it has been shown that ocular torsions occur frequently in patients with strabismus and correlate with the severity of stereovision loss in patients with intermittent exotropia (33). Further studies may test whether treatments of ocular torsion might improve the stereoacuity of those patients.

## 7. Conclusions

The current study showed that the accuracy of depth perception is significantly dependent upon the spatial frequency properties of the observed stimulus when rotating the head. Stereoacuity does not vary when rotating the head around its axis while using narrow-band circle stimuli, but does vary when using conventional broad band random dot tests. Also, the discrepancy between head tilt and ocular-counter roll reported in the literature remarkably does not influence the accuracy of stereovision. We suggest that false dot correspondence account for the loss of stereo sensitivity with head tilt using random dot tests reported in previous studies (e.g. Schreiber et al., 2001) and replicated in the second experiment of the current study. Our results indicate that retinal zones of stereo disparity are not be fixed per se but may be affected by vestibular inputs from off-axis head rotations, which is in agreement with standard energy models of stereo coding and evidence from animal studies. However, this adaptive mechanism may be masked by false correspondences when tested with random dot stimuli. Our findings indicate that patients with abnormal head postures should be tested with sparse as well as broad-band stimuli to evaluate any loss of stereovision.

## Acknowledgment

This research was funded by the National Institutes of Health R01 EY032162. Portions of the current study were presented at the Virtual-Vision Science Society Conference, 2020.

## Authors contributions

PJB developed the study’s overarching research question, designed the protocol of the first experiment.

AC and PJB wrote the code for the psychophysical procedure of first experiment.

AC recruited participants and collected data for the first experiment.

JS, AC, PJB analysed the data for the first experiment.

JS and PJB designed the protocol of the second experiment.

JS recruited participants and collected the data for the second experiment.

JS and PJB analysed the data for the second experiment.

JS wrote the first draft of the manuscript.

All authors refined the draft version to its final submitted form.

